# A Field-Deployable Arsenic Sensor Integrating *Bacillus Megaterium* with CMOS Technology

**DOI:** 10.1101/2024.07.18.604150

**Authors:** Chelsea Y. Hu, John McManus, Fatemeh Aghlmand, Elin Larsson, Azita Emami, Richard M. Murray

## Abstract

Bacteria innately monitor their environment by dynamically regulating gene expression to respond to fluctuating conditions. Through synthetic biology, we can harness this natural capability to design cell-based sensors. *Bacillus megaterium*, a soil bacterium, stands out due to its remarkable heavy metal tolerance and sporulation ability, making it an ideal candidate for heavy metal detection with low transportation costs. However, challenges persist: the synthetic biology toolkit for this strain is underdeveloped and conventional whole-cell sensors necessitate specialized laboratory equipment to read the output. In our study, we genetically modified *B. megaterium* for arsenic detection, establishing a detection threshold below the EPA recommendation of 10 ppb for drinking water in both vegetative cell form and spore form. Additionally, we integrated both engineered *B. megaterium* living cells and spores with CMOS chip for field-deployable arsenic detection. We show that the limit of detection of our integrated sensor is applicable in soil and air arsenic contamination testing. As a proof of concept, this work paves the way for deploying our sensor in resource-limited settings, ensuring real-time arsenic detection in challenging environments.

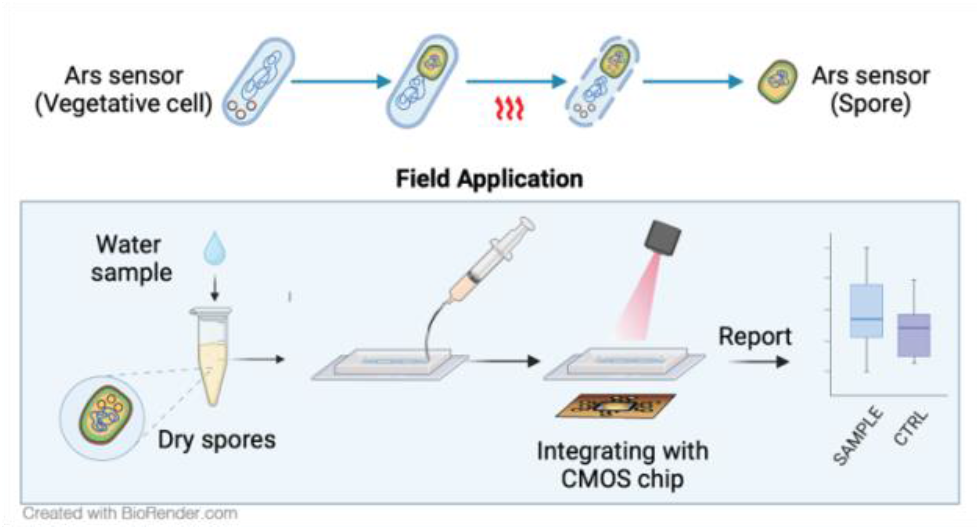

## Introduction

Arsenic is a naturally occurring element that is widely distributed in the Earth’s crust, with significant presence in groundwater, soil, crops, and various minerals. Chronic exposure to arsenic, even at low levels, poses severe health risks including skin lesions, cardiovascular diseases, and various forms of cancer. The US Environmental Protection Agency (EPA) has set the maximum contaminant level for arsenic in drinking water at 10 parts per billion (ppb) or 0.13 micromolar (μM); it also set the soil screening levels (SSLs) for residential areas at 0.39 mg/kg^1^. Occupational Safety and Health Administration (OSHA) has established a permissible exposure limit (PEL) of 10µg/m^3^ over an 8-hour workday, and the US Food & Drug Administration (FDA) has set an action level for rice cereals for infants at 100 ppb. The risks to human health and known permissible arsenic levels demonstrate the importance of monitoring arsenic concentrations in the environment. Traditional methods for arsenic detection, such as atomic absorption spectroscopy (AAS) and inductively coupled plasma mass spectrometry (ICP-MS), offer high sensitivity and accuracy but are often limited by their high cost, complex sample preparation, and requirement for sophisticated laboratory infrastructure. These limitations underscore the need for innovative, cost-effective, and portable solutions for real-time arsenic monitoring in diverse environmental settings.

Whole-cell biosensors, a class of biosensors utilize living cells to detect environmental contaminants. These sensors leverage the natural ability of cells to respond to chemical stimuli, producing a measurable signal in response. *Escherichia coli* is one of the most extensively engineered organisms for whole-cell arsenic sensors, capable of detecting arsenic through the expression of reporter fluorescent or colorimetric proteins in response to arsenic exposure, some reaching low detecting limit below 10ppb in water.^2–5^ Despite their high sensitivity, these biosensors are unsuitable for field applications due to their inability to survive in harsh environments with potential arsenic contamination. Thus, an important consideration when developing a whole-cell biosensor is choosing an appropriate chassis microbe, well-adapted for a specific environment or application.

In the context of environmental monitoring, hardiness is an important factor in whole-cell sensor options. Spores, the metabolically dormant form of certain bacteria, have a high tolerance to a variety of environmental stressors, making them highly suitable for whole cell sensors. Use of spores eliminates the need for cold-chain management, significantly improving the feasibility of whole-cell sensors for environmental monitoring. Several species of the gram-positive genus *Bacillus* are known for producing a type of highly retractile endospores as a survival mechanism.^6^ These endospores germinate and become vegetative cells when the environmental conditions are more favorable. Previous work have shown that engineered whole-cell biosensors can be preserved as spores, for long periods of time.^7,8^ One of the endospore-forming *Bacillus* strains, *B. megaterium*, a ubiquitous soil bacterium, is well-adapted to environments outside the lab, mitigating many biosafety concerns. In addition to its spore-forming capability and environmental ubiquity, *B. megaterium* has several additional traits advantageous for biosensor development: its larger size allows for higher gene expression and a large metabolic load capacity, which is a critical factor for biosensor efficacy.^9,10^ Additionally, it is highly resistant to toxins, including arsenic.^11^ Despite these benefits, *B. megaterium* is relatively under-characterized as a synthetic biology chassis. In this study, we aim to expand its potential as a field-deployable whole cell biosensor by characterizing its sensing capabilities and combining this utility with complementary metal-oxide-semiconductor (CMOS) technology.

CMOS technology has a variety of applications. However, traditional CMOS chips have limited capacity for detecting specific analytes such as arsenic. We recently developed a CMOS chip capable of reading fluorescent signals from living cells, demonstrating the potential of integrating biological and electronic systems.^12^ Additionally, due to its versatility, compact size, and low cost, it is an ideal candidate for portable and field-deployable sensors. The integration of CMOS with whole cell sensors takes advantage of the sensitivity and robustness of living cells with the practicality of CMOS technology, paving the way for robust, field-deployable biosensors.

In this study, we engineered an integrated arsenic sensor for field application by combining whole-cell *B. megaterium* with our customized CMOS chip. This system combines robust characteristics of *B. megaterium* and the portability of the CMOS chip, to achieve field-deployable arsenic detection. To this end, we constructed an arsenic-sensing gene circuit and transformed into *B. megaterium*. We demonstrated the limit of detection (LOD) for arsenic of 0.01 µM and 0.05 µM in vegetative cells and germinated spores, respectively. These detection limits are below the EPA’s maximum contaminant level for arsenic in drinking water of 10 ppb (0.13 µM). In the integrated version of our sensor, we demonstrated a LOD of 0.3 µM and 1.0 µM, for vegetative cells and spores, respectively. These LODs make the system a viable candidate in monitoring inorganic arsenic concentration in soil and air. This integrated approach effectively combines the high sensitivity of biological detection with the practicality of CMOS technology, offering a cost-effective and portable solution for real-time environmental monitoring of arsenic.

## Results and Discussion

### Design and Characterization of whole-cell arsenic biosensor in Bacillus Megaterium

We designed the arsenic sensor utilizing the *ars* operon, which is known for its role in conferring arsenic resistance to *Bacillus*.^13^ Within this operon, the ArsR protein acts as a repressor, binding to the promoter *P*_*ars*_ to inhibit transcription. Exposure to arsenic induces a conformational change in ArsR, leading to its detachment from DNA and thus initiating transcription.^14^ As illustrated in Figure 1a, in our design, the *ArsR* gene is expressed constitutively; the *P*_*ars*_ promoter then regulates the transcription of a superfolder fluorescent protein (sfGFP), which has been long used as the fluorescent reporter in *B. megaterium*. The presence of arsenic activates the sfGFP signal, indicating the presence of the metalloid.

**Figure 1.**
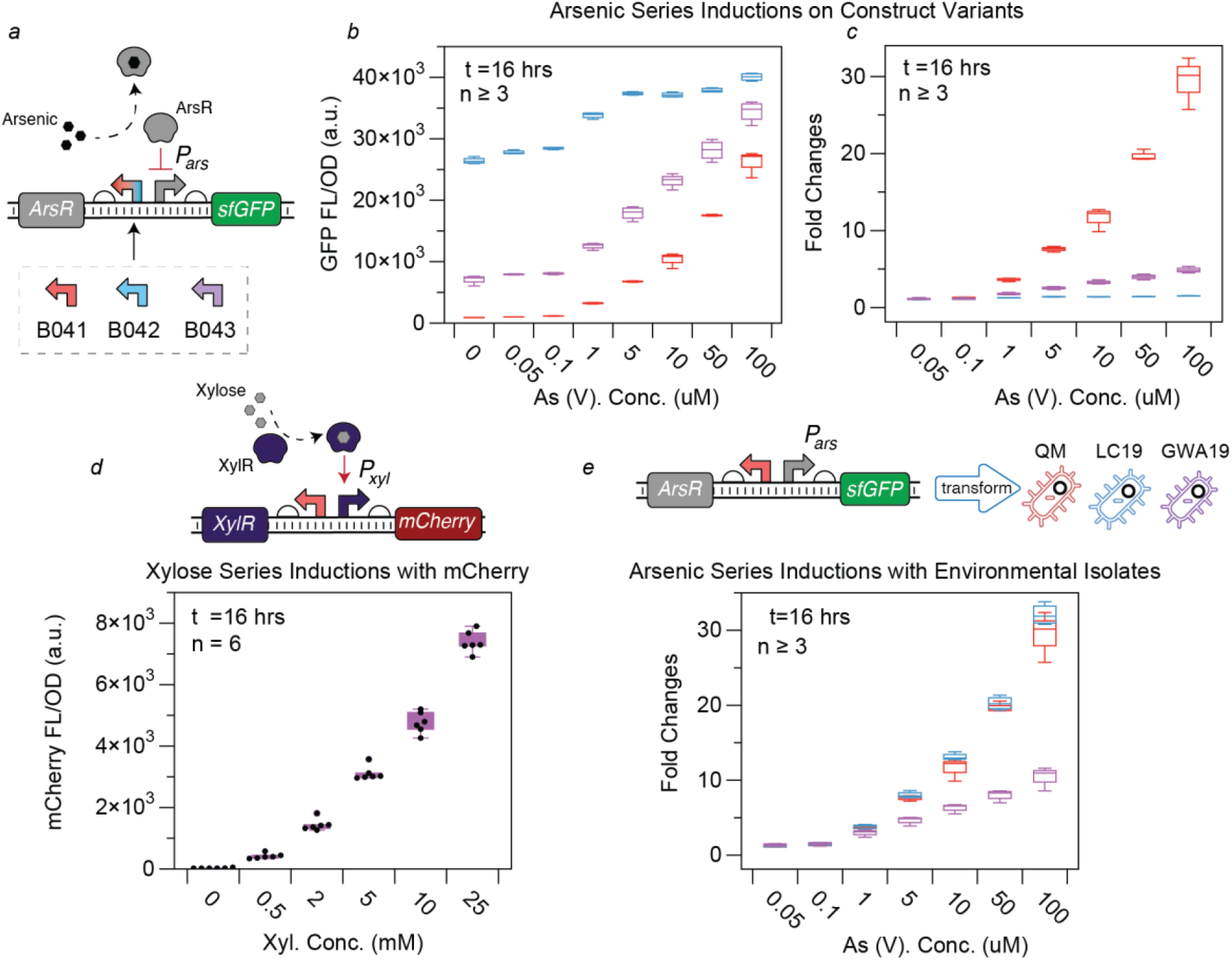
Design and characterization of a whole-cell arsenic sensor in Bacillus Megaterium. (a) Circuit design: the arsenic sensor utilizes the *ars* operon and sfGFP. Transcription regulator ArsR binds to the P_ars_ promoter, repressing downstream transcription. This repression is relieved in the presence of arsenic, and sfGFP is expressed. Three versions of constructs, each with a different constitutive promoter driving ArsR, are characterized. (b) Sensor induction: the sensor was induced with As (V) ranging from 0.05μMto 100μM. The response curves for the three versions of constructs, colored in red (B041), blue (B042), and purple (B043), each with a different constitutive promoter driving ArsR. (c) Normalized response: the same experiment as plotted in (b), with all values normalized to the none-induced conduction (0μM) to highlight fold activation. (d) Characterization of mCherry: The mCherry fluorescent protein was characterized, with induction by xylose concentrations ranging from 0.5 to 25 mM. The constitutively expressed XylR binds to Xylose to activate the expression of mCherry. (e) Arsenic series inductions in environmental isolates: The B041 construct was transformed into two environmental isolates of *B. megaterium*; *B. megaterium* LC19, and *B. megaterium* GWA19, along with the model strain *B. megaterium* QM, As (V) concentration ranged from 0.05μM to 100μM, showing sensor functionality across strains.

We first aim to optimize the fold activation range of the arsenic sensor by varying the expression strength of the transcription regulator ArsR. To do this, we constructed three sensor variants, each utilizing a different constitutive promoter (B041-Pyknw, B042-yngC, and B043-metA), all found in *Bacillus subtilis*.^15^ Through titration experiments (shown in Figures 1b and 1c), we assessed the influence of ArsR promoter strength on sensor functionality. We observed that the construct B042 (blue) that utilizes the weakest promoter for ArsR expression showed significant leaky expression of GFP; the construct B041 (red) that utilizes the strongest promoter for ArsR expression showed minimal leaky expression of GFP (Figure1b, FigureS1a). This is consistent with the mechanistic nature of ArsR functioning as a transcriptional repressor; weak expression of ArsR results in incomplete repression of the *P*_*ars*_ promoter. To evaluate the fold activation, we calculated the ratio of the highest average induced signal to the baseline uninduced signal. The construct with the B041 (red) demonstrated a 30 fold change upon arsenic induction. Based on these findings, we selected promoter P_yknw_ for further development of our arsenic sensor.

Next, we aim to confirm that *B. megaterium* can operate in conditions that are compatible with the CMOS chip. The first component that requires confirmation is the fluorecent protein. Our previous studies have demonstrated robust signal readout from the chip with mCherry,^12^ motivating us to test mCherry’s expression and response dynamics in *B. megaterium*. We utilized the *xyl* operon, previously shown to be functional in *B. megaterium* ^16^ to drive mCherry expression. This system employs a transcriptional regulator xylR that activates the *P*_*xyl*_ promoter in the presence of xylose. Upon induction with xylose, mCherry expression was observed, with the fluorescence signal showing clear differentiation across a range of xylose concentrations from 0 to 25 mM, as detailed in Figure 1d. This result confirms that mCherry can be effectively expressed in *B. megaterium* and respond with a good dynamic range to varying levels of xylose. Additionally, we also tested the activation with both Lysogeny Broth (LB) and M9 media, as plotted in Figure S1b and Figure S1c, we found that although *B. megaterium* is able to grow in the minimal M9 medium with a lower final optical density at 600nm (OD_600_), the *B. megaterium* in minimal M9 did not express any fluorecent protein upon induction. Hence, for the rest of the project, we adopted LB media for both cell culture and sensor induction.

Using a modified transformation procedure (detailed in the Supplementary Information) based on the protocol published by Vorobjeva et al.^17^, we transformed our best-performing construct, B041 (highlighted in red in Figure b and c), into two additional environmental isolates of *B. megaterium*. As shown in Figure 1e, both *B. megaterium* LC 19 and *B. megaterium* GWA19, along with B. *megaterium QM*, were successfully transformed with our plasmid hosted by the pMM1522 backbone. The arsenic sensor was functional in all three strains. However, the sensor transformed into *B. megaterium* GWA19 exhibited a reduced fold activation, indicating strain-to-strain differences in the whole-cell sensor function.

### Characterization of arsenic whole-cell biosensor in vegetative Bacillus Megaterium

Having separately characterized the performance of the arsenic sensor and the CMOS chip-compatible fluorescent protein mCherry, we integrated these components into our final design, as shown in Figure 2a. This design retains the genetic context from B041, using a constitutive P_yknw_ promoter to drive the expression of ArsR, with mCherry replacing sfGFP as the reporter.

First, we assessed the arsenic tolerance of B. *megaterium*. Literature has established that the *Bacillus* species exhibit remarkable resilience to arsenic due to the *ars* operon^13^. We compared the arsenic tolerance of wildtype *B. megaterium QM* with wildtype *B. subtilis* in an As (V) series titration up to 5mM. Our results showed that 2mM and 5mM of As(V) induces 50% and 80% cell death in *B. subtilis*, respectively. In contrast, *B. megaterium* QM did not exhibit detectable cell death even at 5mM of arsenic (V) (Figure S1d, S1e), confirming its ultra-high tolerance to arsenic. Subsequently, we investigated the growth rates of our engineered strain in the presence of As (V). As shown in Figure 2b, we compared the growth curves of the sensor under high concentrations of arsenic (V), ranging from 100 µM to 1 mM, over 16 hours. All cultures were able to grow, despite variations in maximum OD and prolonged lag phases as the concentration increased. Notably, no growth impairment was observed at 100 µM, 200 µM, and 400 µM of As (V) compared to the uninduced culture, suggesting that the presence of As (V) has little impact on the intrinsic metabolism of the *B. megaterium* host chassis at concentrations up to 400 µM.

Next, we conducted a low concentration range titration from 0.05 µM to 100 µM of arsenic. Figure 2c presents a full 16-hour time course of the experiment. After 16 hours, we analyzed the fold changes in fluorescence relative to the uninduced culture across all arsenic concentrations. As depicted in Figure 2d, mCherry expression increased with arsenic concentrations up to 200 µM. Beyond 400 µM, the fluorescence reached a plateau and then started to decline, which aligns with our findings in Figure 2b, where cell growth was impaired starting at 400 µM, indicating that both gene expression and cell growth are impaired at this concentration.

**Figure 2.**
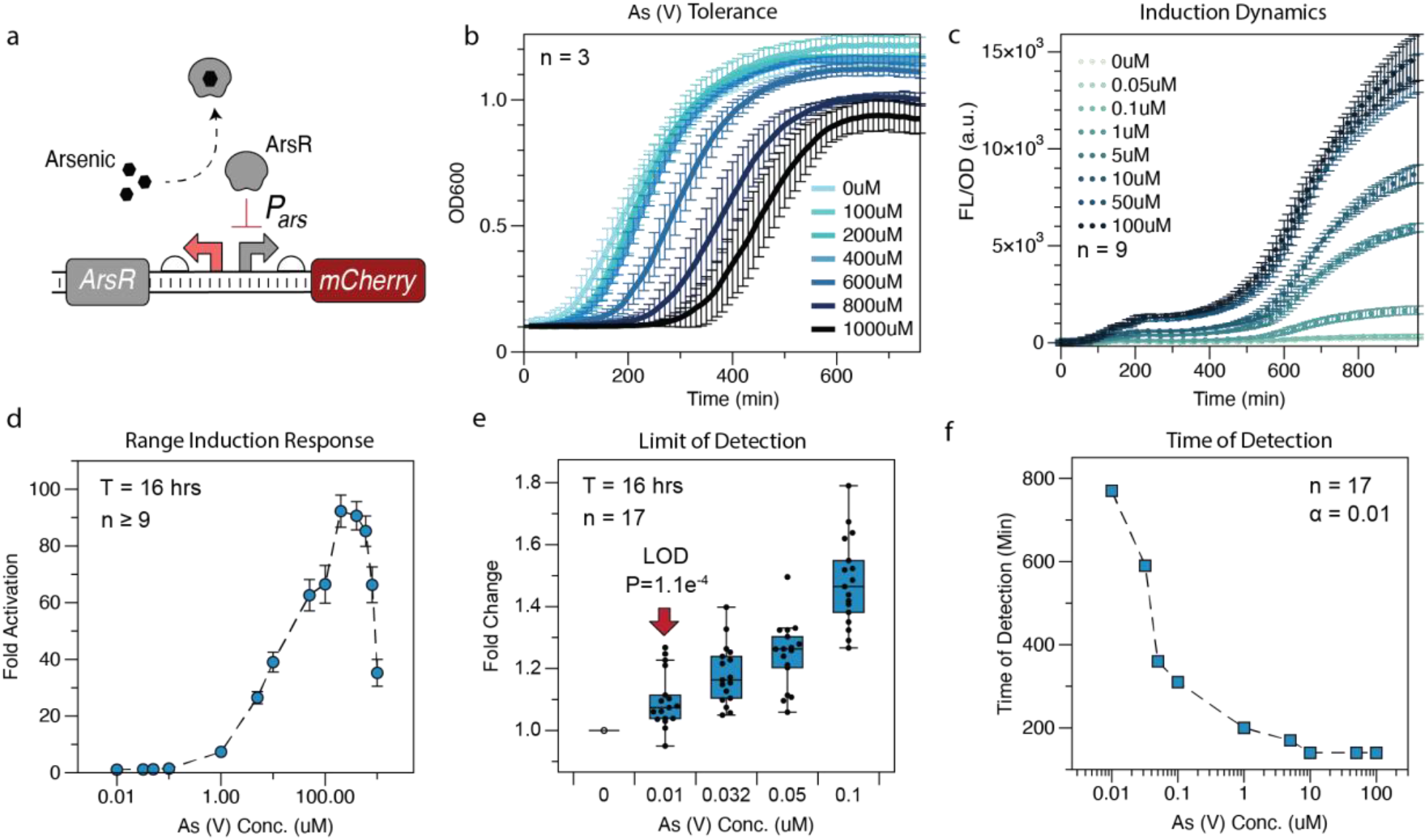
Sensor performance in vegetative B. megaterium. (a) Sensor construct design. The design of the arsenic sensor construct derived from B041, shown in Figure 1a, using mCherry as the reporter (b) Growth dynamic in high arsenic concentrations: Growth dynamics of *B. megaterium* in the presence of As (V). ranging from 100μM to 1000μM, over a 16-hour period. (c) Fluorescent signal dynamics: mCherry fluorescence normalized by OD_600_ over 16 hours, following induction at time =0 min. (d) The sensor response to arsenic concentrations: Sensor response to As (V) ranged from 0.0μM to 1mM, normalized to the 0μM induction culture, measured after 16 hours post-induction. (e) Limit of detection: Induction response of the sensor to low-range As (V) concentrations, spanning from 0.01μM to 0.1μM. The plot shows statistical significance in sensor activation with as low as 0.01μM of As (V). (f) Time of detection. The time required for the whole-cell sensor to register a positive detection of arsenic, ranging from 0.01μM to 100μM, using a two-tailed t-test with a significance threshold of 0.01. The number of biological replicates (n) used in each experiment is indicated in all plots.

To establish the limit of arsenic detection at low concentrations, we explored the detection threshold. Figure 2e shows the fold change in fluorescence of 17 independent colonies exposed to low arsenic doses, ranging from 0.01 µM to 0.1 µM, revealing a detection limit as low as 0.01 µM after 16 hours. In Figure 2f, we plotted the time required for each culture to register a positive detection of arsenic. This was determined by identifying the earliest time point that passed the two-tailed t-test with a significance level threshold of 0.01. The results indicated that detecting 0.01 µM of arsenic takes approximately 770 minutes, while higher concentrations such as 10 µM, 50 µM, and 100 µM are detected much faster, around 140 minutes. Given that the timing does not significantly vary across these higher concentrations, a full 16-hour time course is required to accurately discriminate among them.

### Characterization of arsenic whole-cell biosensor with B. megaterium spores

An advantage of using *Bacillus* strains as chassis for whole-cell biosensors is their ability to sporulate. These spores are metabolically dormant and can therefore survive for long periods of time without nutrients and/or water in extreme environments. Once nutrients are available, these spores can rapidly germinate, becoming metabolically active vegetative cells. This characteristic, in the context of microbial biosensors, prolongs shelf-life, removes cold chain considerations, and widens the scope of deployment for these sensors.^7^ This advantage is the key in designing sensors for point-of-need use, especially in austere environments, where preservation capabilities are minimal.

We first tested the sensor performance using the spores of our engineered *B. megaterium*. As illustrated in Figure 3a, we used a modified procedure based on by Periago et al.^18^ to induce sporulation. Following sporulation, we scraped off the mixture of vegetative cells and spores.

**Figure 3.**
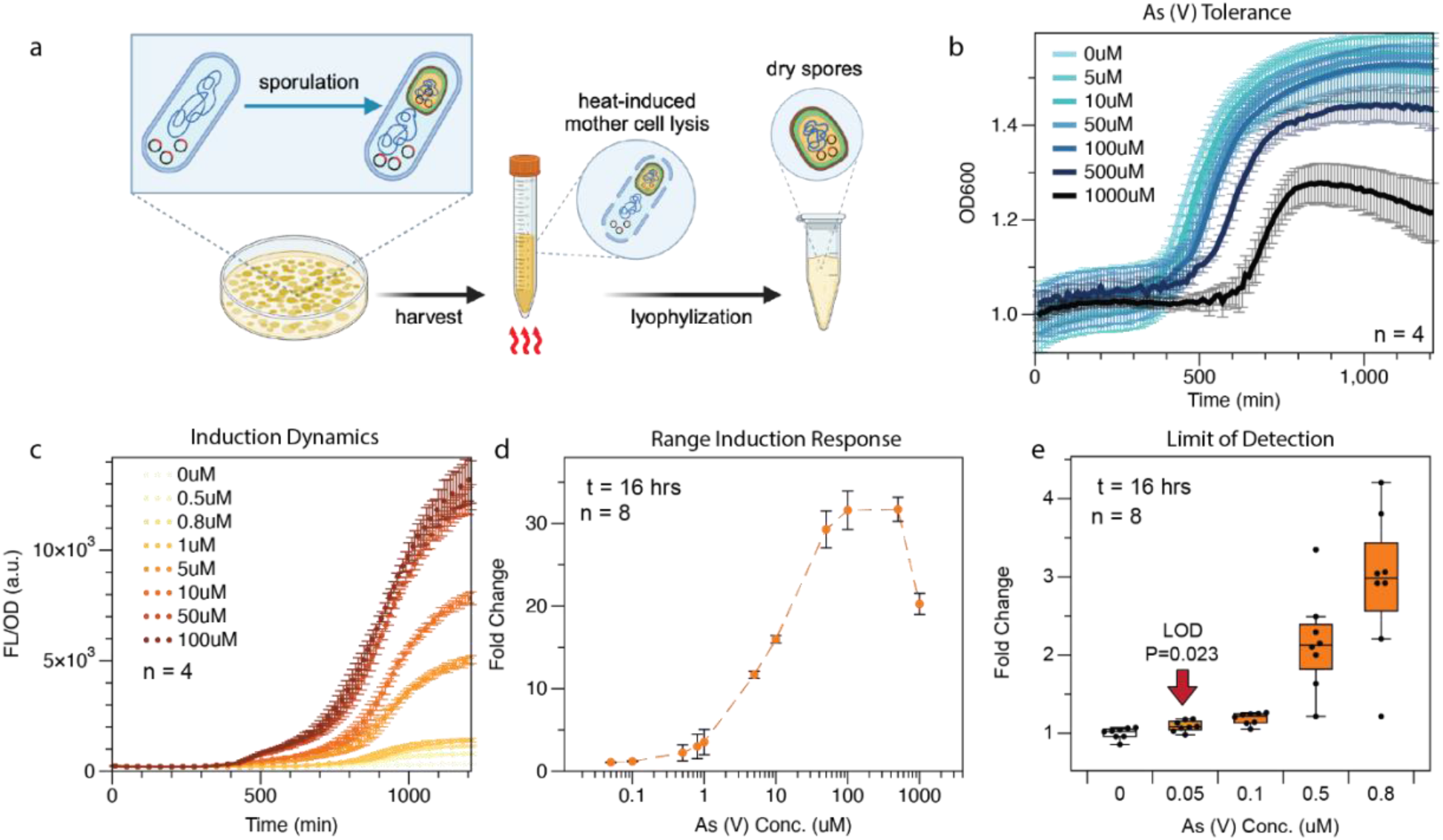
Sensor characterization with spores. (a) Sporulation and spore collection procedure. Illustration created with Biorender.com (b) Sporulation dynamics exposed to high arsenic concentrations ranging from 5μM to 1000μM over 16 hours. Spores were hydrated to OD=1.0 in LB media at t=0. (c) Fluorescent signal dynamics: mCherry fluorescence normalized by OD_600_ over 16 hours, following hydration with LB media containing Arsenic ranging from 0.5μM to 100μM at time =0 min. (d) Sensor response to Arsenic: Spore-based sensor response to As (V) ranged from 0.05μM to 1mM, normalized to the 0μM induction culture, measured 16 hours post-hydration. (e) Limit of detection: Induction response of the sensor to low-range As (V) concentrations, spanning from 0.05μM to 0.8μM. Plot shows statistical significance (p<0.05) in sensor activation with as low as 0.05 μM. The number of technical replicates (n) used in each experiment is indicated in all plots, as spores were harvested in bulk.

This mixture was subjected to high heat to eliminate all vegetative cells, leaving only spores. We divided the spores into two groups: fresh and lyophilized spores, both subjected to arsenic inductions directly without pre-germination.

First, we tested whether *B. megaterium* spores can germinate in arsenic-containing solution and perform sensing simultaneously. We achieve this, we back diluted fresh spores to an OD_600_ value of 1.0 in PBS and hydrated the cultures with water containing a range of arsenic concentrations (0 µM to 1 mM), supplemented with 10X LB media. The growth profile, presented in Figure SI3a, showed significant germination (reaching OD 1.1) at different times based on arsenic concentration: 350 minutes for 0 µM and 5 µM; 420 minutes for 10 µM and 50 µM; 470 minutes for 100 µM; 520 minutes for 500 µM; and nearly 800 minutes for 1 mM. This suggests that in comparison to vegetative cells, spore germination is less resistant to arsenic toxicity. Nevertheless, despite the slight delay in germination, the fluorescence signal increases along with As (V) concentrations, up to 200 µM, indicating relevant sensor performance up to this concentration. We also obtained the induction repone profiles at low concentration of As (V), the detection limit for arsenic was established at 0.05 µM, as shown in Fig S2c.

We repeated the experiment with lyophilized spores stored at room temperature for three weeks. The results, as shown in Figures 3b and 3c, indicated that it took approximately 440 minutes for cultures exposed to 5 µM to 100 µM arsenic to germinate (reach OD 1.1), while for 500 µM arsenic it took 500 minutes, and for 1 mM arsenic it took 660 minutes. Although lyophilized spores showed a slight delay in germination compared to fresh spores, the presence of arsenic up to 100 µM did not significantly impact germination time. After 16 hours, the fluorescence and OD_600_ signals were sufficiently distinct to differentiate concentrations up to 100 µM (Fig 3c), with a maintained LOD at 0.05 µM (Fig 3e).

Our findings affirm that *B. megaterium* spores can function effectively as a durable, long-term viable chassis for biosensors. While the LOD and arsenic tolerance of these spores is subpar of those of vegetative cells, the spores offer substantial advantages in terms of storage flexibility and ease of transportation. Furthermore, lyophilization and storage of the sensor spores do not appear to impair As (V) sensitivity. This trade-off highlights the spores’ potential utility in situations where robustness and portability are more critical than the highest sensitivity or fastest response time.

### Integrating the whole-cell sensor with CMOS for arsenic detection

To realize the potential of point-of-need use of the spores-based sensor, the fluorescent readouts need to be measured without bulky and expensive laboratory instruments. Here, we integrated our biosensor in both *B. megaterium* in the forms of vegetative cells and spores with CMOS chip from our previous work^12^ This 65-nm CMOS chip implements a 600-700 nm range on-chip bandpass optical filter, photodiodes, and processing circuitry, making it capable of reading both the OD_600_ and the mCherry fluorescent protein (Ex/Em: 587nm/610nm) when using a 550nm-LED as the excitation light source.

For performance evaluation, we tested the sensor’s response using vegetative cells as a reference. This setup follows the same procedure as shown in Figure 2, where fresh *B. megaterium* cells were cultured for 16 hours at 37°C post arsenic series induction ranging from 0.032 µM to 100 µM. The mCherry fluorescent response was measured with our CMOS chip in two separate experiments, each with 3-12 biological replicates. As shown in Figure 4b, the integrated sensor demonstrated a LOD of 0.3 µM. This is compared to the response shown in Figure 2e, where a commercial plate reader with excitation at 587 nm and emission at 610 nm achieved an LOD of 0.01 µM. The reduction in sensitivity is significant; however, future advancements in CMOS technology are likely to improve the sensitivity of our system.

**Figure 4.**
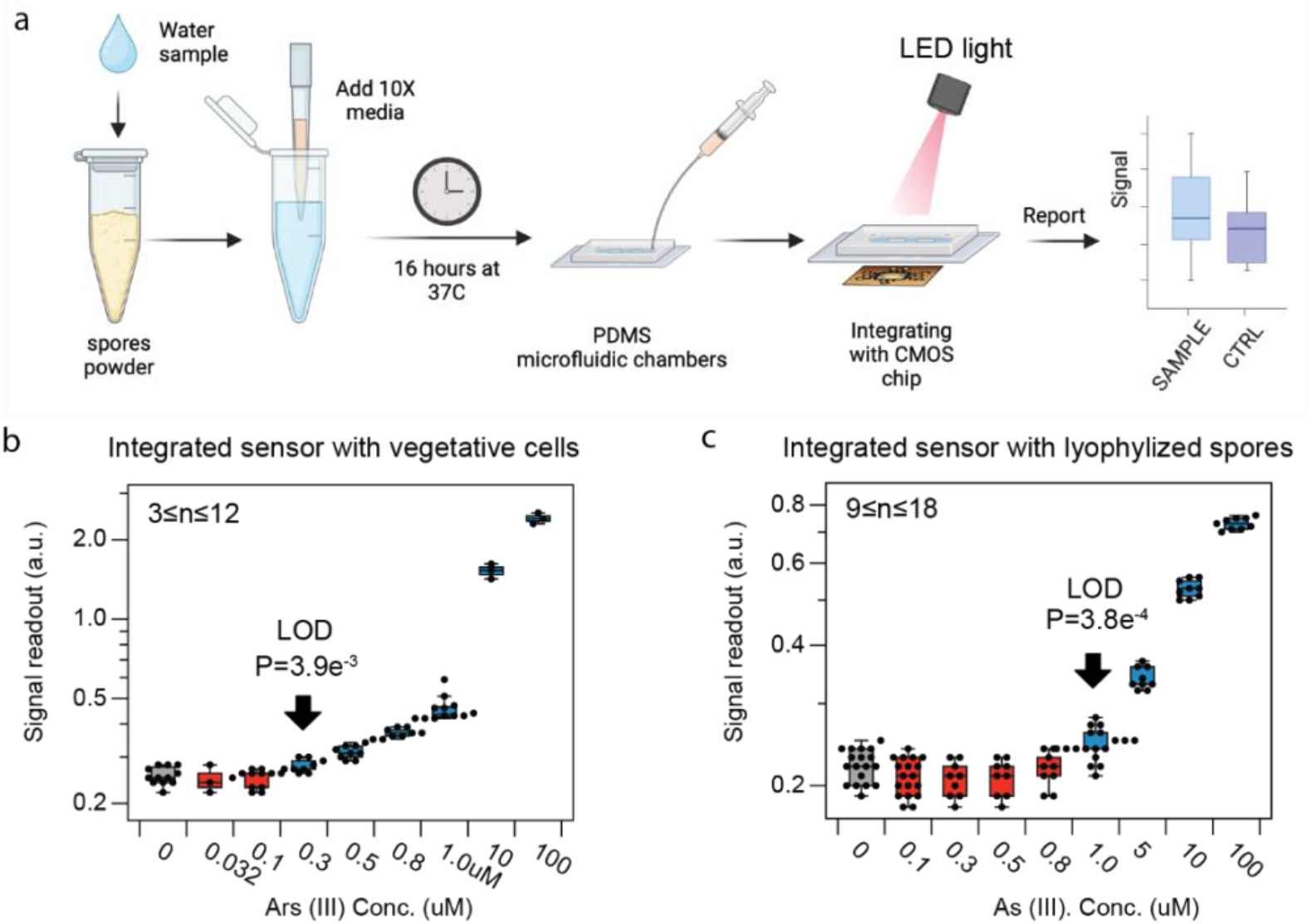
Integrated arsenic sensor characterization. (a) Illustration integrated arsenic sensor detection operation flow, partially created with Biorender.com. (b) Limit of detection when integrating CMOS chip with vegetative cells: Induction response of sensor to As (V) concentrations, ranging from 0.032μM to 100μM. Plot shows statistical significance in sensor activation with as low as 0.3 μM. n represents the number of biological replicates. (c) Limit of detection when integrating CMOS chip with lyophilized spores: Induction response of sensor to As (V) concentrations, ranging from 0.1 μM to 100μM. Plot shows statistical significance in sensor activation with as low as 1.0 μM. n represents technical replicates, as spores were harvested in bulk.

Subsequently, we integrated the CMOS chip with lyophilized spores to simulate a field-testing scenario. As depicted in Figure 4a, we added 900 µL of As (V)-contaminated water sample and 100 µL of 10X LB nutrient into lyophilized spores stored for over six weeks at room temperature. The spores were incubated for 16 hours at 37°C post-hydration before measurement with the CMOS chip. The system was titrated with arsenic from 0.1 µL to 100 µL, in each condition, 9 to18 measurements as technical replicates were obtained. Through two-tailed t-test, we found that the arsenic LOD with this setup is 1.0 µM.

As presented in Table 1, the 5 version of arsenic sensors we have tested are relevant in their corresponding application scenarios. For drinking water, both the EPA and WHO set the acceptable limit for arsenic in drinking water at 10ppb, which corresponds to approximately 0.13μM of arsenic. This makes the sensitivity of our *B. megaterium* sensor in both vegetative cells and spore forms, within application range. For soil arsenic content, the regulation limits vary drastically from country to country. Utilizing established arsenic extraction methods from soil and sediments, such as a combination of phosphate solution and microwave, achieves about 80% arsenic recovery.^19^ This method involves extracting 1 gram of soil into 20mL of phosphate solution,^19^ leading to a LOD of 1.875 mg/kg of arsenic in soil. This is under the guideline limit by 19 states in the U.S. and most international guideline, including Finland at 5mg/kg, Canada at 12mg/kg and UK at 32mg/kg, etc.^1^ In theory, the sensitivity of our integrated sensor (CMOS chip + lyophilized spores) is applicable in testing arsenic content in soil using current inorganic arsenic extraction methods by most international standards and some US state standards. To achieve EPA’s SSLs for residential areas at 0.39 mg/kg,^1^ the sensitivity of our integrated sensor needs to be further improved. To test arsenic content in rice, various extraction methods achieve a recovery efficiency ranging from 75% to 100%.^20^ Using a setup of 1g of rice and 20mL of extracting solution, to meet the FDA’s recommendation of 0.1 mg/kg of inorganic arsenic in rice cereals for infants,^21^ the sensor’s LOD needs to be as low as 0.05 μM at a 75% recovery rate. In air monitoring, (OSHA) has set the permissible exposure limit (PEL) for inorganic arsenic at 10ug/m^3^ in air, averaged over an 8-hour workday. Using OSHA’s midget air bubbler sampling method, which bubbles up to 120 liters of air into 10 milliliters of solution at a rate of 1 liter per minute.^22^ To sample air that contains 10ug/m^3^ pf arsenic, it takes 75 minutes to sample 75 liters of air to achieve solution concentration of 1μM, which agrees with the sampler’s setup. This means that the integrated sensor is theoretically applicable for field testing of airborne arsenic using a standard midget air bubbler.

**Table 1.** Limit of detection of all versions of arsenic sensor and their potential applicable scenarios.

| Whole-cell sensor type | Electronics | LOD in $\mu\text{M}$ (p value) | Figure | Water<sup>1</sup> | Soil<sup>2</sup> | Rice<sup>3</sup> | Air<sup>4</sup> |
| --- | --- | --- | --- | --- | --- | --- | --- |
| Vegetative cells | Plate reader | 0.01 $\mu\text{M}$ (1.1e-4) | Figure 2 | ● | ● | ● | ● |
| Fresh Spores | Plate reader | 0.05 $\mu\text{M}$ (0.025) | Figure S2 | ● | ● | ● | ● |
| Lyophilized spores | Plate reader | 0.05 $\mu\text{M}$ (0.023) | Figure 3 | ● | ● | ● | ● |
| Vegetative cells | CMOS chip | 0.3 $\mu\text{M}$ (3.9e-3) | Figure 4b | ✗ | ◎ | ✗ | ● |
| Lyophilized spores | CMOS chip | 1.0 $\mu\text{M}$ (3.8e-4) | Figure c | ✗ | ◎ | ✗ | ● |

1. Based on EPA's standard for drinking water, at 10 ppb. 2. Based on EPA's SSL for residential areas at 0.39mg/kg (solid circle) and most international standards between 5-15mg/kg (hollow circle). 3. Based on FDA's arsenic limit at 0.1mg/kg in rice cereals for infants. 4. Based on OSHA's PEL of 10ug/m<sup>3</sup> of arsenic in air. ● applicable, ✗not applicable, ◎ partially applicable.

## Conclusion

In this work, we successfully demonstrated the potential of *B. megaterium* as a chassis for developing a whole-cell biosensors. By harnessing the natural resilience and sporulation capabilities of *B. megaterium*, we created a sensitive and portable arsenic detection system that integrates with CMOS technology. The whole-cell sensor performed effectively in both vegetative cell and spore forms, with detection limits of 0.01 µM and 0.05 µM, respectively, well below the EPA’s arsenic limit for drinking water. The integration with CMOS chips allows for portable and field-deployable applications, providing a significant reduction in equipment size and cost compared to traditional methods. The CMOS-based sensor maintained a detection limit of 0.3 µM with vegetative cells and 1.0 µM with spores, suitable for various environmental monitoring scenarios, including soil and air arsenic contamination.

This work highlights the advantages of using *B. megaterium* spores for long-term viability and ease of transport, making the sensor highly applicable for resource-limited settings. The successful integration of biological and electronic systems in this study paves the way for deploying robust, cost-effective, and real-time environmental monitoring solutions. Future enhancements in the sensitivity of the integrated sensor could broaden its applicability, ensuring compliance with stringent regulatory standards for arsenic detection in diverse environments.

Such improvements could be achieved by enhancing the sensitivity of the CMOS technology, which includes increasing the active area of photodetectors and optical filters to collect more photons/signals, redesigning the on-chip optical filters to better match the target wavelengths, and increasing the bias current of the front-end to reduce noise at the expense of slightly higher power consumption. Additional future work includes conducting field tests and calibrating the sensor with various samplers to standardize the procedure for different applications.

## Materials and Methods

### Construction of Arsenic Sensors

The *arsR* gene and *pArs, yknW, aprE, metA*, and *yngC* promoters, derived from *B. subilitis*^23^, were synthesized by IDT. The pMM1522 vector (Mobitech) was linearized by PCR-amplification (Forward primer: 5’ cattactcgcatccattctcaggctgtctcgtctcgtctcatgcgcaaaccaacc 3’, Reverse primer: 5’ gcttggattctgcgtttgtttccgtctacgaactcccagcttaagtgaacgcaaaggtta 3’). Parts were assembled, along with the sfGFP gene, using overlapping UNSs^24^ and NEBuilder (New England Bio Labs). Cloning was carried out in *E. coli* DH10B cells. Plasmids were purified using a Qiagen QIAprep Spin Miniprep Kit (Qiagen 27104). Linear fragments were gel extracted and purified using MinElute Gel Extraction Kit (Qiagen 28606). Part sequences and plasmid maps are provided in the Supporting Information (Tables S1 and S2). Confirmed DNA plasmids were transformed into *B. megaterium* protoplasts based on method previously established^25^, detailed protocol is also included in the Supporting Information.

### Bulk Fluorescent Experiments

Bulk fluorescence measurement was employed to evaluate the expression of fluorescent proteins following chemical induction. For vegetative cells, overnight liquid cultures were initiated from freshly isolated colonies on LB Agar plates, and then incubated in LB medium at 37°C with shaking at 220 rpm for 16 hours. Subsequently, a 200 µL subculture was established in a 96-well culture block by combining 196 µL of LB medium with antibiotics and 4 µL of the overnight culture. This subculture was grown for 4 hours, then diluted 20-fold with LB medium containing the appropriate antibiotics (TetR) and arsenic inducers in Na_2_HAsO_4_ · 7H_2_O (CAS 10048-95-0). The diluted cultures were incubated at in a 96-well optic plate using a BioTek Synergy H1 plate reader for 16 hours. The plate reader maintains conditions at 37 °C with maximum linear shaking, measuring fluorescence (sfGFP-Ex/Em: 485nm/528nm or mCherry-Ex/Em: 587nm/610nm) and OD_600_ every 10 minutes. For lyophilized spores, a mixture of 180 µL of arsenic-containing deionized water and 20 µL of 10X LB medium was used to rehydrate the spores to an OD of 1.0. These rehydrated spores were then grown in a 96-well plate in the BioTek Synergy H1 plate reader for 24 hours under the same conditions, with measurements of fluorescence (mCherry-Ex/Em: 587nm/610nm) and optical density OD_600_ taken every 10 minutes.

### As(V) tolerance Assay

*B. subtilis* and *B. megaterium* tolerance to As(V) were assessed on a dose-response curve. Overnight cultures were diluted to OD_600_= 0.01 and were grown until OD_600_= 0.6; split into 2mL aliquots and exposed to a range of As(V) concentrations (final concentrations: 0mM, 0.5mM, 1mM, 2mM, 5mM) for 16h with shaking at 37°C. After exposure, 1mL of each concentration was harvested via centrifugation, washed with PBS (Fisher; BP2944100) with a final resuspension of 0.5mL. Viability was analyzed following manufacturer instructions using *Bac*Light Bacterial Viability Kit (Invitrogen; L7012). Survival was determined by a change in normalized fluorescence compared to 0mM As (V) exposure using BioTek Synergy H1). Confocal microscopy was performed by immobilizing samples on 2% agarose pads (Fisher; BP1356-500) using SP8 (Leica). Images were acquired and processed with Leica Application Suite software. Fluorescence was measured for SYTO9 (Ex/Em: 480nm/500nm) and Propidium Iodide (Ex/Em: 490nm /635nm). The sensitivity assay had three independent experiments in triplicate performed.

### Preparation of Spores from Bacillus megaterium

Cells were initially cultured from a single colony in 5 mL of LB medium at 37°C, agitated at 220 rpm overnight. Following this, 200 µL aliquots were evenly distributed onto spore-forming media plates, prepared with 5 g tryptone, 2.5 g yeast extract, 1 g glucose, 15 g agar, and deionized water up to 1 L, with the pH adjusted to 7.0, with appropriate antibiotics. These plates were then incubated at 30°C for a duration of 5 days. Subsequently, the grown biomass was scrapped of from the plates and re-suspended in deionized water in 15 mL conical tubes, with the suspension diluted to achieve an OD_600_ of 1.0. To ensure the inactivation of vegetative cells, the tubes containing the suspension were placed in a water bath at 80°C for 10 minutes, then allowed to cool at 20°C for another 10 minutes. The final step involved lyophilizing the suspension within the same 15 mL conical tubes, preparing it for further analysis or storage. Lyophilized spores were stored in room temperature.

### Statistical Analysis

Fluorescence per optical density (FL/OD) signals were adjusted by subtracting the background, which was determined using the average autofluorescence signal derived from the FL/OD measurements of three colonies of non-transformed cells. Data analysis was conducted using Microsoft Excel. The sample size was not predetermined through statistical methods. All single colonies were selected randomly from agar plates following the transformation of cells with purified circular DNA, employing antibiotics as selection markers. No manual methods for group allocation were implemented. Each plate originated from a distinct transformation event, and all colonies on a plate were considered biological replicates. Throughout the experiments, no data were excluded from analysis.

The LOD was determined statistically. For vegetative cells, we calculated the signal fold changes for each colony by dividing the fluorescent output of an induced culture by the fluorescent output of the uninduced culture, to account for autofluorescence. For spores, the signal fold changes for each measurement were calculated by dividing the fluorescent output of an induced culture by the average fluorescent output of the uninduced culture. The sensor’s LOD was established when the fold changes at a given concentration were significantly higher than those of the uninduced group, based on a two-tailed t-test with P<0.05. For CMOS experiments, signals were analyzed without fold activation calculation, but the same statistical tests and significance thresholds were used to determine the LOD.

## Author Contribution

RMM, AE, JM, and CYH conceptualized the project, CYH and JM designed the experiments, CYH, JM, FA, and EL conducted the experiments, CYH performed data analysis and wrote the manuscript.

## Acknowledgement

The authors would like to thank Tracy Mei at Texas A&M University for their contributions on the arsenic tolerance assay and the manuscript. This research is partially supported by the Institute for Collaborative Biotechnologies (ICB) through contract W911NF-19-D-0001 from the U.S. Army Research Office. In addition to the support from ICB, the study also received support from the Caltech Center for Sensing to Intelligence (S2I) and Heritage Medical Research Institute. The content of the information does not necessarily reflect the position or the policy of the government, and no official endorsement should be inferred.

## References

1. Teaf, C. M., Covert, D. J., Teaf, P. A., Page, E. & Starks, M. J. Arsenic Cleanup Criteria for Soils in the US and Abroad: Comparing Guidelines and Understanding Inconsistencies. in vol. 15 (2010).

2. Baumann, B. & Meer, J. R. van der. Analysis of Bioavailable Arsenic in Rice with Whole Cell Living Bioreporter Bacteria. J. Agric. Food Chem. 55, 2115–2120 (2007).

3. Jia, X., Bu, R., Zhao, T. & Wu, K. Sensitive and Specific Whole-Cell Biosensor for Arsenic Detection. Appl. Environ. Microbiol. 85, (2019).

4. Yoon, Y., Kim, S., Chae, Y.Jeong, S.-W. & An, Y.-J. Evaluation of bioavailable arsenic and remediation performance using a whole-cell bioreporter. Sci. Total Environ. 547, 125–131 (2016).

5. Huang, C.-W.Wei, C.-C. & Liao, V. H.-C. A low cost color-based bacterial biosensor for measuring arsenic in groundwater. Chemosphere 141, 44–49 (2015).

6. McKenney, P. T., Driks, A. & Eichenberger, P. The Bacillus subtilis endospore: assembly and functions of the multilayered coat. Nat. Rev. Microbiol. 11, 33–44 (2013).

7. Date, A., Pasini, P., Sangal, A. & Daunert, S. Packaging Sensing Cells in Spores for Long-Term Preservation of Sensors: A Tool for Biomedical and Environmental Analysis. Anal. Chem. 82, 6098–6103 (2010).

8. Valenzuela-García, L. I. et al. Design of a Whole-Cell Biosensor Based on Bacillus subtilis Spores and the Green Fluorescent Protein To Monitor Arsenic. Microbiol. Spectr. 11, e00432–23 (2023).

9. Kumari, W. M. N. H., Thiruchittampalam, S., Weerasinghe, M. S. S., Chandrasekharan, N. V. & Wijayarathna, C. D. Characterization of a Bacillus megaterium strain with metal bioremediation potential and in silico discovery of novel cadmium binding motifs in the regulator, CadC. Appl. Microbiol. Biotechnol. 105, 2573–2586 (2021).

10. Stammen, S. et al. High-Yield Intra- and Extracellular Protein Production Using Bacillus megaterium. Appl. Environ. Microbiol. 76, 4037–4046 (2010).

11. Njoku, K. L., Akinyede, O. R. & Obidi, O. F. Microbial Remediation of Heavy Metals Contaminated Media by Bacillus megaterium and Rhizopus stolonifer. Sci. Afr. 10, e00545 (2020).

12. Aghlmand, F. et al. A 65-nm CMOS Fluorescence Sensor for Dynamic Monitoring of Living Cells. IEEE J. Solid-State Circuits 58, 3003–3019 (2023).

13. Sato, T. & Kobayashi, Y. The ars Operon in the skin Element of Bacillus subtilis Confers Resistance to Arsenate and Arsenite. J. Bacteriol. 180, 1655–1661 (1998).

14. Wu, J. & Rosen, B. P. Metalloregulated expression of the ars operon. J. Biol. Chem. 268, 52–58 (1993).

15. Song, Y. et al. Promoter Screening from Bacillus subtilis in Various Conditions Hunting for Synthetic Biology and Industrial Applications. PLoS ONE 11, e0158447 (2016).

16. Stammen, S. et al. High-Yield Intra- and Extracellular Protein Production Using Bacillus megaterium. Appl. Environ. Microbiol. 76, 4037–4046 (2010).

17. Vorobjeva, I. P., Khmel, I. A. & Alföldi, I. Transformation of Bacillus Megaterium Protoplasts by Plasmid DNA. FEMS Microbiol. Lett. 7, 261–263 (1980).

18. Periago, P. M., Delgado, egoña, Fernández, P. S. & Palop, A. Bacillus megaterium Spore Germination and Growth Inhibition by a Treatment Combining Heat with Natural Antimicrobials. Food Technol. Biotechnol 1, 23 (2005).

19. Yuan, C.-G., He, B., Gao, E.-L., Lü, J.-X. & Jiang, G.-B. Evaluation of extraction methods for arsenic speciation in polluted soil and rotten ore by HPLC-HG-AFS analysis. Microchim. Acta 159, 175–182 (2007).

20. Costa, B. E. S., Coelho, L. M., Araújo, C. S. T., Rezende, H. C. & Coelho, N. M. M. Analytical Strategies for the Determination of Arsenic in Rice. J. Chem. 2016, 1–11 (2016).

21. U.S. Food and Drug Administration. Supporting document inorganic arsenic in Rice cereals action levels. https://www.fda.gov/food/chemical-metals-natural-toxins-pesticides-guidance-documents-regulations/supporting-document-action-level-inorganic-arsenic-rice-cereals-infants.

22. Occupational Safety and Health Administration. OSHA Technical Manual (OTM) - Section II: Chapter 1 https://www.osha.gov/otm/section-2-health-hazards/chapter-1#sampling_plan.

23. Song, Y. et al. Promoter Screening from Bacillus subtilis in Various Conditions Hunting for Synthetic Biology and Industrial Applications. PLoS ONE 11, e0158447 (2016).

24. Torella, J. P. et al. Unique nucleotide sequence–guided assembly of repetitive DNA parts for synthetic biology applications. Nat. Protoc. 9, 2075–2089 (2014).

25. Biedendieck, R. et al. Chapter ten Systems Biology of Recombinant Protein Production Using Bacillus megaterium. Methods Enzym. 500, 165–195 (2011).

