## Supporting Information for "A Field-Deployable Arsenic Sensor Integrating *Bacillus Megaterium* with CMOS Technology"

Chelsea Y. Hu[1,3], John McManus[1], Fatemeh Aghlmand[1], Elin Larsson[1], Azita Emami[2], and Richard M. Murray[1]

1. Division of Biology and Bioengineering, California Institute of Technology, Pasadena, CA.
2. Division of Engineering and Applied Science, California Institute of Technology, Pasadena, CA.
3. Department of Chemical Engineering, Texas A&M University, College Station, TX

### I. Supplemental experimental results

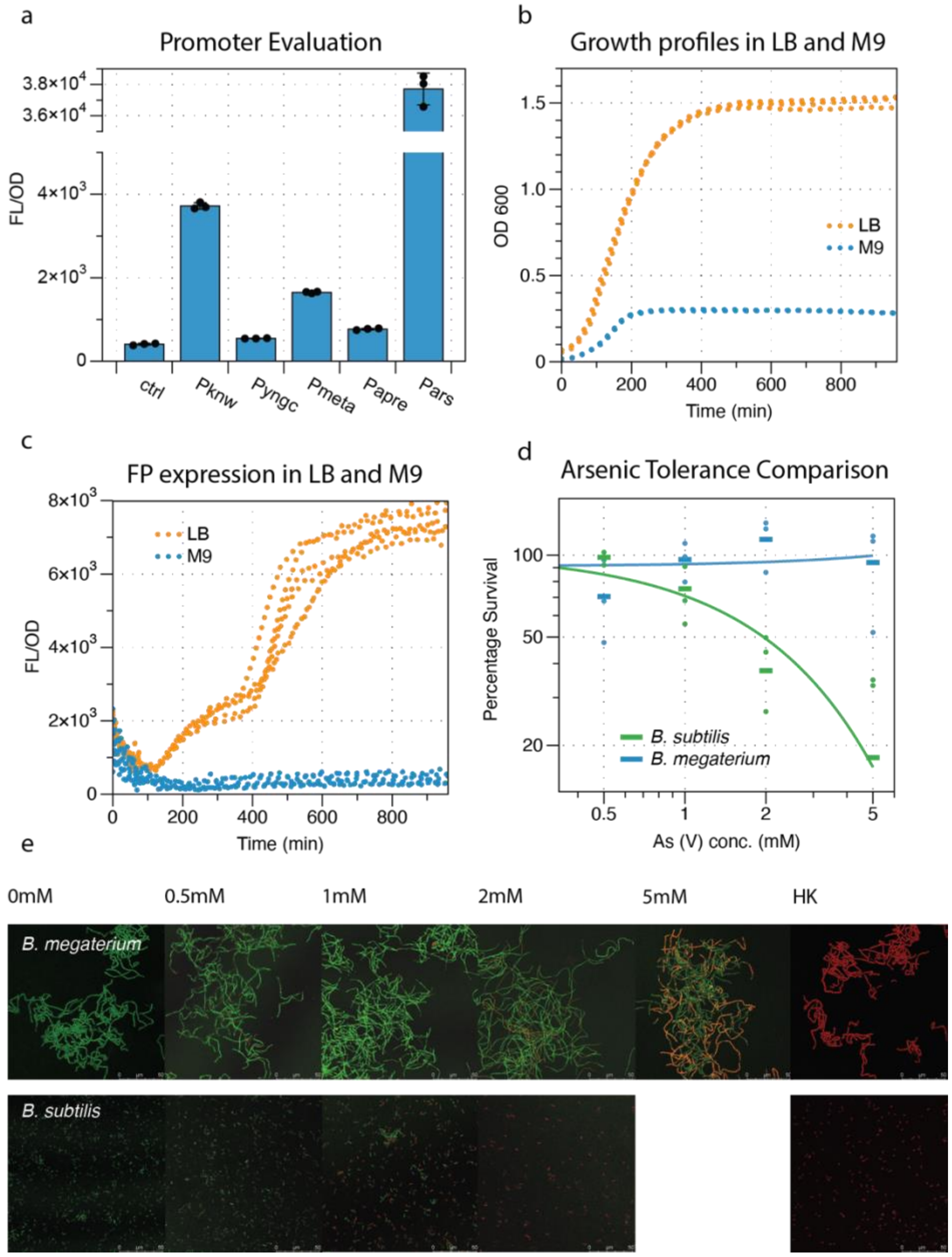

***Figure S1 Constitutive promoters, growth conditions, and arsenic tolerance.*** *(a) Relative promoter strengths characterization in B. megaterium. Each promoter drives the transcription of GFP. (b) Growth dynamics of B. megaterium over time when culture grown in LB and M9 media. (c) GFP expression dynamics over time when cultured in LB and M9 medium. (d) Viability of B. megaterium (blue) and B. subtilis (green) after exposure to As(V). Plotted with 3 biological replicates (dots) and their averages (bars).(e) Microscopy images of B. megaterium after As(V) exposure and labeled with SYTO9 and propidium Iodide overlayed to differentiate live (green) from dead (red) bacteria. HK=Heat Killed.*

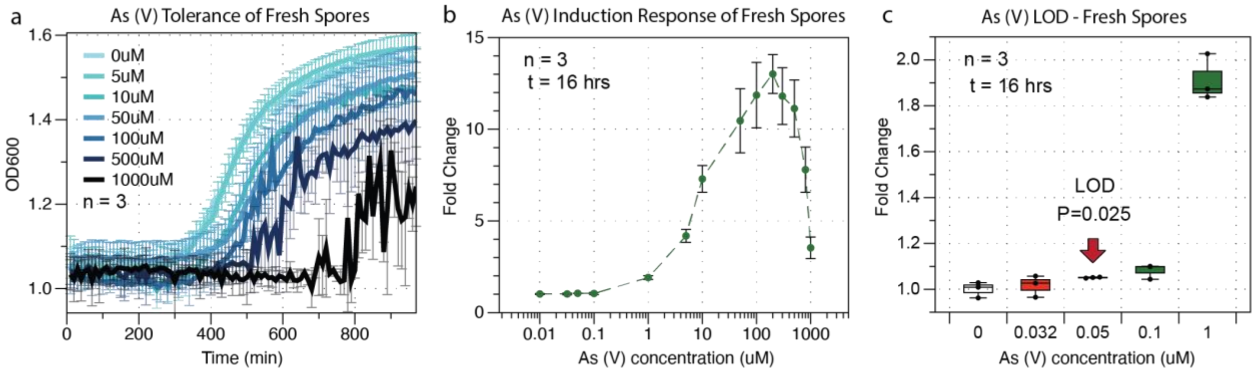


***Figure S1*** *Sensor characterization with fresh spores. (a) Sporulation dynamics exposed to high arsenic concentrations ranging from 5uM to 1000uM over 16 hours. Spores were hydrated to OD=1.0 in LB media at t=0. (b) Sensor response to arsenic: Spore-based sensor response to As (V) ranged from 0.05uM to 1mM, normalized to the 0uM induction culture, measured 16 hours post-hydration. (c) Limit of detection: Induction response of sensor to low range As (V) concentrations, ranging from 0.032uM to 1uM. Plot shows statistical significance in sensor activation with as low as 0.05 uM. n denotes for the number of technical replicates.*

### *II.* Protoplast Transformation of *Bacillus Megaterium*

**For 8 transformations, scale accordingly:**

**1. Initiate Culture**: Inoculate a 5mL LB medium with bacteria from a fresh plate. Incubate at 37°C with continuous shaking at 220rpm overnight.
**2. Scale-Up Culture**: Transfer 1mL of the overnight culture to 50mL of fresh LB medium in a baffled flask. Grow at 37°C with shaking until an optical density (OD) of 1.0 is reached. This usually takes about 2 hours.
**3. Cell Pelleting**: Centrifuge the culture in a 50mL Falcon tube at 2600g for 15 minutes at 4°C to pellet the cells.
**4. Lysozyme Treatment**: Resuspend the cell pellet in 5mL of SMMP solution with 2mg of lysozyme added. Secure the tube on a rocker, ensuring alignment with the rocking axis, and incubate at 37°C for 30 minutes. From this step onward, handle samples gently to avoid mechanical stress.
**5. Protoplast Harvesting**: Centrifuge at 1300g for 10 minutes at room temperature (RT), then carefully pour off the supernatant.
**6. Wash to Remove Lysozyme**: Resuspend the protoplasts in 5mL of SMMP and centrifuge again (1300g for 10 minutes at RT) to pellet. Carefully remove the supernatant.
**7. Repeat Washing**: Perform the wash step described in step 6 once more to ensure thorough removal of lysozyme.
**8. Protoplast Resuspension**: Resuspend the washed protoplasts in 4mL of SMMP.
**9. Plasmid Addition**: For each transformation, mix 500µL of the protoplast suspension with up to 100µL of DNA solution (totaling 5µg of DNA) in a new 50mL Falcon tube.
**10. PEG-P Addition**: Add 1.5mL of PEG-P to each transformation mix. Incubate at room temperature for 2 minutes.
**11. Dilution and Mixing**: Add 5mL of SMMP to the mixture and gently roll the tube to mix.
**12. PEG Removal**: Centrifuge at 1300g and room temperature for 10 minutes to pellet the protoplasts. Decant the supernatant with care.
**13. Protoplast Rescue**: Add 500µL of SMMP to each tube to resuspend the protoplasts.
**14. Standing Incubation**: Incubate the tubes at 30°C for 45 minutes without shaking.
**15. Shaking Incubation**: Continue incubation at 30°C for another 45 minutes, this time with slow shaking at approximately 50rpm.
**16. Plating**: Gently mix the protoplasts with 2.5mL of CR5 top agar. Immediately before plating, mix CR-b (kept at 50°C) into the CR5 top agar (at room temperature). Ensure the agar is cool but not solidifying. Maintain the mixture at 37°C if needed to prevent solidification before plating on LB agar plates containing the appropriate antibiotics.
**17. Incubation for Colonization**: Incubate the plated cultures at 30°C for 24 hours to allow for colony formation.

### Buffers and Media

| | | | |
|---|---|---|---|
| ***2x AB3 (1L)*** *– autoclave 15 min*<br>3 g beef Extract<br>3 g yeast Extract<br>10 g peptone<br>1g glucose<br>7 g NaCl<br>7.36 g $K_2HPO_4$<br>2.64 g $KH_2PO_4$ | ***2x SMM (1L)****- filter sterilize*<br><br>1 M Sucrose<br>40 mM maleic acid<br>40 mM MgCl<br>pH 6.5 | ***CR-a (250 mL)****- filter sterilize*<br><br>51.5 g sucrose<br>3.25 g MOPS<br>0.33 g NaOH<br>pH 6.5 | ***CR-5 salts****- autoclave sterilize*<br><br>1.25g $K_2SO_4$<br>50g $MgCl_26H_2O$<br>0.25g $KH_2PO_4$<br>11g $Cacl_2xH_20$<br>Add 625ml $H_2O$ |
| ***casamino acid sln****-* filter sterilize<br><br>10g casamino acid<br>100 mL MiliQ | ***PEG-P*** *(100ml)*<br><br>40g PEG in 40ml water, autoclave for 15 mins, add 36mL sterilized 2XSMM buffer. Total volume should be ~100ml | ***12% proline*** *- filter sterilize*<br><br>12g in 100ml miliQ | ***20% glucose*** *- filter sterilize*<br><br>20g in 100ml miliQ |
| ***CR-b (14.25 mL)*** *– need 2.5ml/ transformation, autoclave for 20 min,* ***make day of, keep at 50C after.***<br>200 mg agar<br>500 mg yeast extract<br>10mg casamino acid (100uL of 100g/L solution) | ***SMMP*** *- for 8 transformations*<br><br>25ml 2xAB3<br>25ml 2XSMM | ***Lysozyme stock***<br><br>100mg/ml in water, made in small volume, filter sterilize, then store in -20C.<br><br>Use 20uL for 2mg | ***CR5 top Agar –*** made for one transformation, scale accordingly.<br><br>2.5ml CR-a<br>575uL CR5 salts<br>250 uL 12% proline<br>250 ul 20% glucose<br>1.42ml CR-b (this is agar, add right before plating) |

### Important Reminder:

Following the 30-minute lysozyme digestion period, it is critical to handle the protoplasts with utmost care. When adding any liquid to the samples, ensure it is done gently, allowing the liquid to flow slowly along the inner side of the tube. To mix, use only a gentle rolling motion or slow rocking; avoid shaking or pipetting actions. In cases where pellets are present, allocate a maximum of 5 minutes for careful mixing. It is not necessary to achieve complete suspension.

### III. *DNA sequences involved in this study*

***Table S1. Parts for Sensor Assembly.*** *UNS are underlined.*

pMM1522 Backbone

5’cattactcgcatccattctcaggctgtctcgtctcgtctcatgcgcaaaccaacccttggcagaacatatccatcgcgtccgccatctccagcagccgcacgcggc
gcatctcgggccgcgttgctggcgtttttccataggctccgcccccctgacgagcatcacaaaaatcgacgctcaagtcagaggtggcgaaacccgacaggactat
aaagataccaggcgtttccccctggaagctccctcgtgcgctctcctgttccgaccctgccgcttaccggatacctgtccgcctttctcccttcgggaagcgtggcgctt
tctcatagctcacgctgtaggtatctcagttcggtgtaggtcgttcgctccaagctgggctgtgtgcacgaaccccccgttcagcccgaccgctgcgccttatccggta
actatcgtcttgagtccaacccggtaagacacgacttatcgccactggcagcagccactggtaacaggattagcagagcgaggtatgtaggcggtgctacagagttc
ttgaagtggtggcctaactacggctacactagaaggacagtatttggtatctgcgctctgctgaagccagttaccttcggaaaaagagttggtagctcttgatccggca
aacaaaccaccgctggtagcggtggtttttttgtttgcaagcagcagattacgcgcagaaaaaaaggatctcaagaagatcctttgatcttttctacggggtctgacgct

cagtggaacgaaaactcacgttaagggattttggtcatgagattatcaaaaaggatcttcacctagatccttttaaattaaaaatgaagttttaaatcaatctaaagtatatat
gagtaaacttggtctgacagttaccaatgcttaatcagtgaggcacctatctcagcgatctgtctatttcgttcatccatagttgcctgactccccgtcgtgtagataactac
gatacgggagggcttaccatctggccccagtgctgcaatgataccgcgagacccacgctcaccggctccagatttatcagcaataaaccagccagccggaagggc
cgagcgcagaagtggtcctgcaactttatccgcctccatccagtctattaattgttgccgggaagctagagtaagtagttcgccagttaatagtttgcgcaacgttgttgc
cattgctgcaggcatcgtggtgtcacgctcgtcgtttggtatggcttcattcagctccggttcccaacgatcaaggcgagttacatgatcccccatgttgtgcaaaaaag
cggttagctccttcggtcctccgatcgttgtcagaagtaagttggccgcagtgttatcactcatggttatggcagcactgcataattctcttactgtcatgccatccgtaag
atgcttttctgtgactggtgagtactcaaccaagtcattctgagaatagtgtatgcggcgaccgagttgctcttgcccggcgtcaacacgggataataccgcgccacat
agcagaactttaaaagtgctcatcattggaaaacgttcttcggggcgaaaactctcaaggatcttaccgctgttgagatccagttcgatgtaacccactcgtgcacccaa
ctgatcttcagcatcttttactttcaccagcgtttctgggtgagcaaaaacaggaaggcaaaatgccgcaaaaaagggaataagggcgacacggaaatgttgaatact
catactcttcctttttcaatattattgaagcatttatcagggttattgtctcatgagcggatacatatttgaatgtatttagaaaaataaacaaataggggttccgcgcacatttc
cccgaaaagtgccacctgacgtctaagaaaccattattatcatgacattaacctataaaaataggcgtatcacgaggccctttcgtcttcaagaattcctgttataaaaaa
aggatcaattttgaactctctcccaaagttgatcccttaacgatttagaaatccctttgagaatgtttatatacattcaaggtaaccagccaactaatgacaatgattcctga
aaaaagtaataacaaattactatacagataagttgactgatcaacttccataggtaacaacctttgatcaagtaagggtatggataataaaccacctacaattgcaatacc
tgttccctctgataaaaagctggtaaagttaagcaaactcattccagcaccagcttcctgctgtttcaagctacttgaaacaattgttgatataactgttttggtgaacgaaa
gcccacctaaaacaaatacgattataattgtcatgaaccatgatgttgtttctaaaagaaaggaagcagttaaaaagctaacagaaagaaatgtaactccgatgtttaac
acgtataaaggacctcttctatcaacaagtatcccaccaatgtagccgaaaataatgacactcattgttccagggaaaataattacacttccgatttcggcagtacttagc
tggtgaacatctttcatcatataaggaaccatagagacaaaccctgctactgttccaaatataattcccccacaaagaactccaatcataaaaggtatatttttccctaatc
cgggatcaacaaaaggatctgttactttcctgatatgttttacaaatatcaggaatgacagcacgctaacgataagaaaagaaatgctatatgatgttgtaaacaacataa
aaaatacaatgcctacagacattagtataattcctttgatatcaaaatgaccttttatccttacttctttctttaataatttcataagaaacggaacagtgataattgttatcatag
gaatgagtagaagataggaccaatgaatataatgggctatcattccaccaatcgctggaccgactccttctcccatggctactatcgatccaataagaccaaatgcttta
cccctattttcctttggaatatagcgcgcaactacaaccattacgagtgctggaaatgcagctgcaccagccccttgaataaaacgagccataataagtaaggaaaag
aaagaatggccaacaaacccaattaccgacccgaaacaatttattataattccaaataggagtaaccttttgatgcctaattgatcagatagctttccatatacagctgttc
caatggaaaaggttaacataaaggctgtgttcacccagtttgtactcgcaggtggtttattaaaatcatttgcaatatcaggtaatgagacgttcaaaaccatttcatttaat
acgctaaaaaaagataaaatgcaaagccaaattaaaatttggttgtgtcgtaaattcgattgtgaataggatgtattcacatttcaccctccaataatgagggcagacgta
gtttatagggttaatgatacgcttccctcttttaattgaaccctgttacattcattacacttcataattaattcctcctaaacttgattaaaacattttaccacatataaactaagttt
taaattcagtatttcatcacttatacaacaatatggcccgtttgttgaactactctttaataaaataatttttccgttcccaattccacattgcaataatagaaaatccatcttcat
cggctttttcgtcatcatctgtatgaatcaaatcgccttcttctgtgtcatcaaggtttaatttttatgtatttcttttaacaaaccaccataggagattaaccttttacggtgtaa
accttcctccaaatcagacaaacgtttcaaattcttttcttcatcatcggtcataaaatccgtatcctttacaggatattttgcagtttcgtcaattgccgattgtatatccgattt
atatttatttttcggtcgaatcatttgaacttttacatttggatcatagtctaatttcattgcctttttccaaaattgaatccattgtttttgattcacgtagttttctgtattcttaaaata
agttggttccacacataccaatacatgcatgtgctgattataagaattatctttattatttattgtcacttccgttgcacgcataaaaccaacaagattttattaatttttttatatt
gcatcattcggcgaaatccttgagccatatctgacaaactcttatttaattcttcgccatcataaacatttttaactgttaatgtgagaaacaaccaacgaactgttggctttt
gtttaataacttcagcaacaaccttttgtgactgaatgccatgtttcattgctctcctccagttgcacattggacaaagcctggatttacaaaaccacactcgatacaacttt
ctttcgcctgtttcacgattttgtttatactctaatatttcagcacaatcttttactctttcagcctttttaaattcaagaatatgcagaagttcaaagtaatcaacattagcgatttt
cttttctctccatggtctcacttttccactttttgtcttgtccactaaaacccttgatttttcatctgaataaatgctactattaggacacataatattaaaagaaaccc ccatctat
ttagttatttgtttggtcacttataactttaacagatggggtttttctgtgcaaccaattttaagggttttcaatactttaaaacacatacataccaacacttcaacgcacctttca
gcaactaaaataaaaatgacgttatttctatatgtatcaagataagaaagaacaagttcaaaaccatcaaaaaaagacaccttttcaggtgctttttttattttataaactcatt
ccctgatctcgacttcgttcttttttacctctcggttatgagttagttcaaattcgttctttttaggttctaaatcgtgtttttcttggaattgtgctgttttatcctttaccttgtctac
aaaccccttaaaaacgtttttaaaggcttttaagccgtctgtacgttccttaaggaattctcatgtttgacagcttatcatcgataagctttaatgcggtagtttatcacagtta
aattgctaacgcagtcaggcaccgtgtatgaaatctaacaatgcgctcatcgtcatcctcggcaccgtcaccctggatgctgtaggcataggcttggttatgccggtac
tgccgggcctcttgcgggatatcgtccattccgacagcatcgccagtcactatggcgtgctgctagcgctatatgcgttgatgcaatttctatgcgcacccgttctcgga
gcactgtccgaccgctttggccgccgcccagtcctgctcgcttcgctacttggagccactatcgactacgcgatcatggcgaccacacccgtcctgtggatctgacgc
gtgtaactgcgaaggtaaaacgggtgattgaaaataaactaacaaatgaagaggaacgaaagagaagtctagaatttgttacgtttatatcggatgaattactgcaaaa
tgatacggagcctcgcacatttattattcagaactctcttttatcgctagagaaaatccatactctcgcacagaccaaagaggtagaggaatataaagaagtcataacta
ctctgacaagacatgattcataatttgaaaagcaggtaaactaacccaaaaggcaagactctccgtagtttagaaaagagcaggttgctaataacatataaacagcca
gttgccgttatgataggtgactggctgcatgggatgaaaaggtgagggtggagacagacataacactcttaatagaagagggtaattctttctcttttatagaaaatcaat
taattgaaagtagctccttcattcttaagatcaacgtgatataggtttgctaacctttgcgttcacttaagctgggagttcgtagacggaaacaaacgcagaatccaagc3
’

sfGFP Part

5’gcactgaaggtcctcaatcgcactggaaacatcaaggtcgctgtcaccggatgtgctttccggtctgatgagtccgtgaggacgaaacagcctctacaaataatttt
gtttaatctagagaaagaggagaaatactagatggagaatttatacttccagagtagcaaaggtgaagaactgtttaccggcgttgtgccgattctggtggaactggat
ggcgatgtgaacggtcacaaattcagcgtgcgtggtgaaggtgaaggcgatgccacgattggcaaactgacgctgaaatttatctgcaccaccggcaaactgccgg
tgccgtggccgacgctggtgaccaccctgacctatggcgttcagtgttttagtcgctatccggatcacatgaaacgtcacgatttctttaaatctgcaatgccggaaggc

tatgtgcaggaacgtacgattagctttaaagatgatggcaaatataaaacgcgcgccgttgtgaaatttgaaggcgataccctggtgaaccgcattgaactgaaaggcacggattttaaagaagatggcaatatcctgggccataaactggaatacaactttaatagccataatgtttatattacggcggataaacagaaaaatggcatcaaagcgaattttaccgttcgccataacgttgaagatggcagtgtgcagctggcagatcattatcagcagaataccccgattggtgatggtccggtgctgctgccggataatcattatctgagcacgcagaccgttctgtctaaagatccgaacgaaaaacgggaccacatggttctgcacgaatatgtgaatgcggcaggtattacgtggagccatccgcagttcgaaaaataacattactcgcatccattctcaggctgtctcgtctcgtctc3’

pArs-ArsR Part

5’gctgggagttcgtagacggaaacaaacgcagaatccaagcctccgttgctgtagtagcaaagtagcgatggaggaatataaaaatggaaaacctctatctggttacacaagcttttagcgggttttacaattaatcaaaataaattgatttatttgcttgcattaatttaaaaatcatgagtataataaatacatcaaaaaaaattgattaaagaaggcgatatgaatggatgagacgaaatcagaactgctacggaaatatgaacaaaaatttaaggctcttgctgatcagaaacgattagagatcatgtatgaactttgtcaaagggggaaaaacctgtgtttgtgatctgactgagatttttgaggtgacgcaatctaaactttcttatcatctaaaaattttattggatgccaacttaataacgaaagagacaaagggaacatggagttattatgatctaaatgatgaggaagtaaatggtttattatcagaagagctttgttgtatatttagaaaaaaaggtgaaggagattgctgctaagcactgaaggtcctcaatcgcactggaaacatcaaggtcg3’

pArs Part

5’ctgacctcctgccagcaatagtaagacaacacgcaaagtcctccgttgctgtagtagcaaagtagcgatggaggaatataaaaatggaaaacctctatctggttacacaagcttttagcgggttttacaattaatcaaaataaattgatttatttgcttgcattaatttaaaaatcatgagtataataaatacatcaaaaaaaattgcactgaaggtcctcaatcgcactggaaacatcaaggtcg 3’

ArsR Part

5’gctgggagttcgtagacggaaacaaacgcagaatccaagcttagcagcaatctccttcaccttttttctaaatatacaacaaagctcttctgataataaaccatttacttcctcatcatttagatcataataactccatgttccctttgtctctttcgttattaagttggcatccaataaaatttttagatgataagaaagtttagattgcgtcacctcaaaaatctcagtcagatcacaaacacaggtttttccccttttgacaaagttcatacatgatctctaatcgtttctgatcagcaagagccttaaatttttgttcatatttccgtagcagttctgatttcgtctcatccattcatatcgccttctttaatcaattttttttgatgtagagccaactccctttacaacctcactcaagtccgttagag 3’

yknW Part

5’gagccaactccctttacaacctcactcaagtccgttagagtaacaaatagtacgggaaggatatcaaaaagtttcatgtttttctaataaatttccaaaaacctctgacctcctgccagcaatagtaagacaacacgcaaagtc 3’

yngC Part

5’gagccaactccctttacaacctcactcaagtccgttagagtaatctttttcactattatatgactgattgacgagcaagatggttacaatttttttaccgcactgacctcctgccagcaatagtaagacaacacgcaaagtc 3’

metA Part

5’gagccaactccctttacaacctcactcaagtccgttagaggtttaattttcttcattataggaattttcacgccaattttcaatctatctatacgtgttctgacctcctgccagcaatagtaagacaacacgcaaagtc 3’

aprE Part

5’gagccaactccctttacaacctcactcaagtccgttagagctattctgtgaatttattgtaatagatggaataatattttagtagacccatttttttgagactgacctcctgccagcaatagtaagacaacacgcaaagtc 3’
